# Endothelin-1 signaling regulates chamber-specific mouse atrial cardiomyocyte cytokinesis and polyploidy

**DOI:** 10.64898/2026.05.26.727795

**Authors:** Tian Lan, Sabrina Kaminsky, Lorna Rinck, Yannik Andrasch, Eva Zickgraf, Mahak Singhal, Chi-Chung Wu

## Abstract

Cardiomyocyte (CM) maturation is a central process in postnatal heart development accompanied by profound structural, metabolic, and cell cycle changes. One hallmark of this maturation program is CM polyploidy that is closely associated with the loss of cardiac regenerative capacity. Most insights into ploidy regulation have come from studies of ventricular CMs, whereas the spatiotemporal dynamics and molecular regulation of atrial CM polyploidy remain poorly understood.

We show that CM polyploidy in the postnatal mouse heart is highly chamber-specific, with >90% of ventricular CMs are polyploid, compared with ∼30% of left atrial (LA) CMs and ∼15% of right atrial (RA) CMs. These chamber-specific differences correlate with their differential susceptibility to cytokinesis failure and are regulated, at least in part, by endocardial cells. Mechanistically, we identify the endocardial/endothelial-derived factor EDN1 as a postnatally enriched signal in the LA compared with the RA. EDN1 can act directly on primary aCMs to inhibit cytokinesis, in part by suppressing Wnt signaling. Consistently, inhibition of Edn1 signaling *in vivo* using Bosentan reduced CM cytokinesis failure specifically in the LA. Altogether, our findings reveal a previously unrecognized role for endothelial-myocardial crosstalk in regulating chamber-specific CM polyploidy through Edn1 signaling.

## Introduction

Cardiomyocyte (CM) polyploidization, i.e., multiplication of the genome, is prevalent across mammals^1^, including humans^2,3^. This process is typically the result of cell cycle variants in CMs, where defective karyokinesis results in nuclear polyploidy and cytokinesis failure results in multinucleation^4^, respectively. Although the physiological role of CM polyploidy remains largely unclear, recent studies have established an inverse correlation between the abundance of polyploid CMs and the endogenous regenerative capacity of the heart. In mice, most CMs transition from a diploid to polyploid state during late embryonic to early postnatal development^5^, coinciding with CM cell cycle exit and the loss of regenerative capacity at postnatal day (P) 7^6^. In zebrafish, in which most adult CMs are mononuclear diploid, experimental induction of CM polyploidy significantly impairs CM proliferation and heart regeneration^7^. Consistently, inbred mouse strains with a higher proportion of mononuclear diploid adult CMs exhibit improved functional recovery after myocardial infarction (MI)^8^. Altogether, these findings highlight ploidy as an important factor determining the endogenous regenerative capacity of CMs. In addition to its role during postnatal heart development, CM polyploidization also occurs under pathological conditions. Following MI, the vast majority of cycling adult mouse CMs fail to complete cytokinesis and become multinucleated^9,10^, while in humans, CM nuclear ploidy is significantly increased in patients with heart failure^11^. These observations lead to the hypothesis that re-activation of CM polyploidization may contribute, at least in part, to the limited regenerative capacity of the diseased heart. In support of this idea, experimental strategies that promote successful cytokinesis in adult mouse CMs improve left ventricular systolic function after MI^12^. Hence, understanding how CM ploidy is regulated may provide important insights into postnatal heart development and inform regenerative strategies to replenish CMs after cardiac injury. Most insights into CM polyploidy have emerged from studies of ventricular CMs (vCMs). In mice, vCMs rapidly become multinucleated and reach near adult level within the first two postnatal weeks^5^. In humans, most vCMs become polyploid between 20 to 40 years of age, comprising both multinucleated CMs and mononuclear CMs with polyploid nuclei^2,3,13^. Intriguingly, atrial CMs (aCMs) also become polyploid in mice^14^ and humans^15^, albeit to a lower extent than vCMs. Importantly, aCM polyploidy may also be dynamically regulated under pathological conditions. In patients with atrial fibrillation and heart failure, nuclear ploidy is specifically increased in left atrial (LA) CMs^15^, whereas increased DNA synthesis, multinucleation, and DNA content have been observed in rat aCMs following MI^16,17^. These observations suggest that aCM polyploidization is part of both postnatal maturation and myocardial remodeling in disease. However, the spatiotemporal dynamics, chamber specificity, and molecular regulation of aCM polyploidy remain largely unexplored.

To date, mechanistic investigations of CM polyploidization have largely focused on CM-intrinsic regulators^18^. Apart from fibroblasts^19^ and the extracellular matrix^20-22^, little is known about how non-myocytes influence CM ploidy. Yet, intercellular communication between CMs and non-myocytes is essential for heart development, homeostasis, and disease. In mice, endothelial/endocardial cells (ECs/EdCs) represent the most abundant non-myocyte populations in the heart^23^. Increasing evidence suggest that cardiac ECs/EdCs are not only passive suppliers of oxygen and nutrients, but also serve an instructive role in regulating heart morphogenesis^24,25^, CM proliferation and maturation^26^, and regeneration^27,28^ through angiocrine signals. However, whether endothelial-myocardial crosstalk regulates postnatal CM polyploidization, and which angiocrine signals may be involved, remains unknown.

Here, we show that CM polyploidization in the postnatal mouse heart is highly chamber-specific, with distinct timing and extent across the ventricle, LA, and right atrium (RA). We further identify cardiac ECs/EdCs as regulators of aCM cytokinesis and polyploidy, at least in part through chamber-specific endothelin-1 (Edn1) signaling. Mechanistically, EDN1 promotes aCM cytokinesis failure by suppressing canonical Wnt signaling. Altogether, these findings highlight a previously unrecognized role for endothelial-myocardial crosstalk in CM ploidy regulation during postnatal heart development.

## Results

### Postnatal mouse cardiomyocyte polyploidy is highly chamber-specific

Although the progression of vCM polyploidization has been well-characterized^5^, the spatiotemporal dynamics of aCM polyploidy in the postnatal mouse heart remain unclear. To address this question, we first quantified mouse CM nucleation in single cell suspensions collected at different postnatal stages. Consistent with previous reports^5,14,29^, vCMs rapidly became binucleated between P4 and P9, increasing from ∼25% to ∼80%. In contrast, aCMs, analyzed as pooled left and right atria, showed only a modest increase in multinucleation over the same period (Fig. 1A). We next asked whether polyploidization differs between the two atria by quantifying CM nucleation separately in the LA and RA. Unlike vCMs, which show little chamber-specific variation in multinucleation^14^, LA CMs displayed a significantly higher proportion of binucleation than RA CMs at P7. This difference became even more pronounced by 8 weeks of age, reaching ∼25% in the LA compared to ∼12% in the RA (Fig.1B). Since karyokinesis failure or G2/M arrest can result in mononuclear CMs with polyploid nuclei, we re-analyzed the mononuclear CM population from the samples shown in Figure 1B. We quantified nuclear ploidy based on relative DAPI intensity, using presumably diploid nuclei from non-myocytes as an internal reference. Notably, the vast majority of mononuclear LA and RA CMs had diploid nuclei at all timepoints examined (>90%; Fig. 1C), indicating that karyokinesis failure or G2/M arrest is not a major contributor to aCM polyploidy. To more directly compare the extent of cytokinesis failure between cycling LA and RA CMs, we performed an EdU pulse-chase assay. Mouse pups received a single EdU injection to label cycling CMs, and their nucleation status was assessed after 48 h, corresponding to approximately one cell cycle, as a readout of cytokinesis failure. Consistent with the binucleation trajectory observed in Fig. 1B, a higher proportion of LA CMs underwent cytokinesis failure compared with RA CMs between P3 and P7, whereas no significant difference was observed at P0 (Fig. 1D). When both multinuclear and mononuclear polyploid CMs were considered, ∼30% of LA CMs and ∼15% of RA CMs were polyploid at 8 weeks of age (Fig. 1E). Altogether, these data indicate that CM polyploidization in postnatal mice is highly chamber-specific in both timing and extent, which can be explained, at least in part, by differential susceptibility to cytokinesis failure.

**Figure 1.**
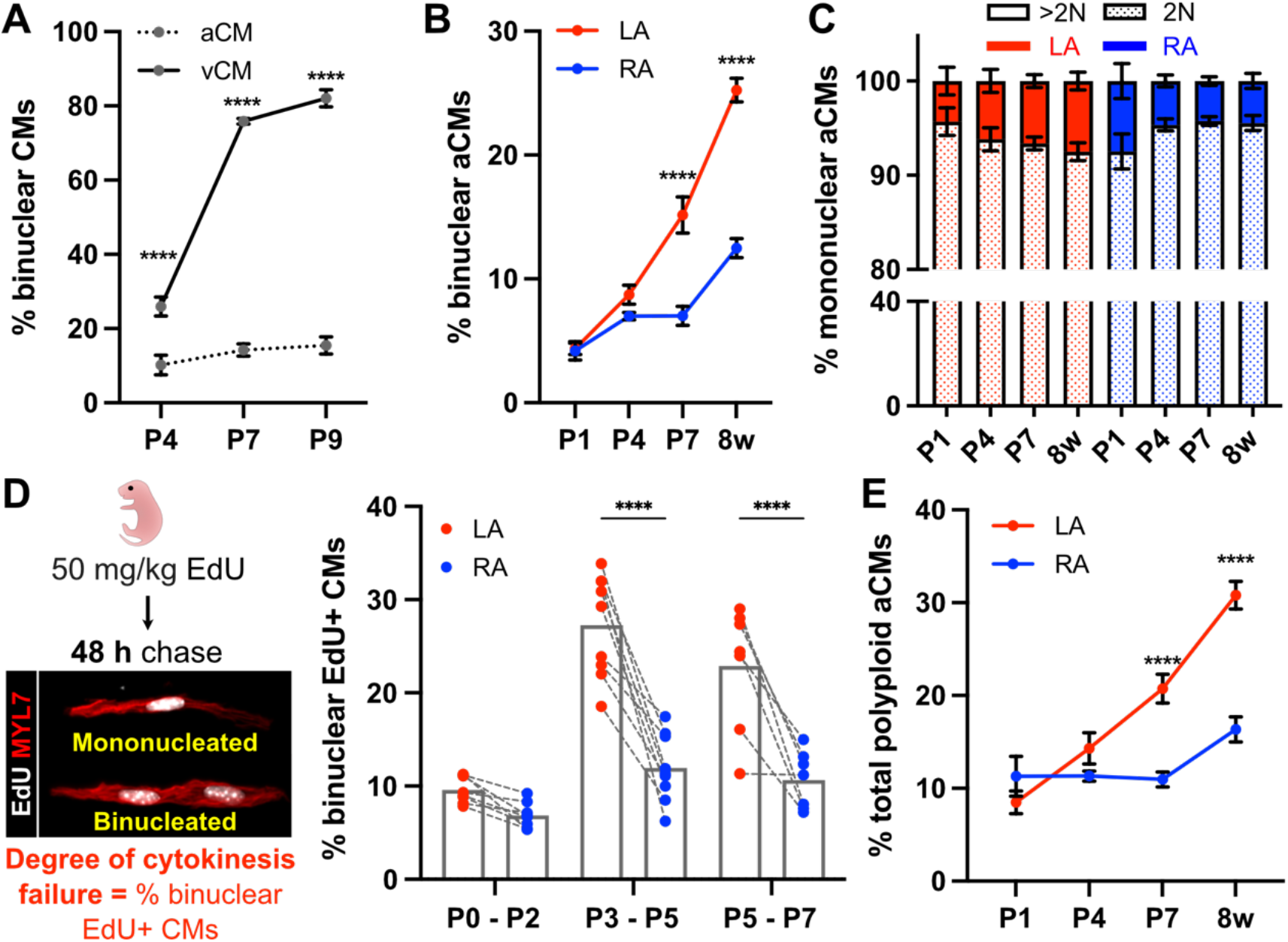
Postnatal mouse CM polyploidization is chamber-specific. **(A)** Quantification of the percentage of binuclear aCMs and vCMs from P4 (n = 6), P7 (n = 5), and P9 (n = 8) mice. **(B)** Quantification of the percentage of binuclear LA and RA CMs from P1 (n = 8), P4 (n = 9), P7 (n = 17), and 8 weeks (n = 9) mice. **(C)** Quantification of nuclear ploidy of the mononuclear CM population from *B*. **(D)** Schematics of the EdU pulse-chase binucleation assay shown on the left. Quantification of the percentage of EdU+ LA and RA CMs shown on the right; P0 to P2 (n = 8), P3 to P5 (n = 9), P5 to P7 (n = 7). LA and RA samples from the same mouse were connected with dashed line. **(E)** Percentage of total polyploid CMs (mononuclear polyploid + binucleated) calculated from *B* and *C*. Values represent mean ± s.e.m. *A, B, D, E* (* *P < 0*.*0332, \**\*\* *P < 0*.*0002*, **** *P < 0*.*0001* by Two-Way ANOVA followed by Šidak test); *C* (*P > 0*.*05* by Two-Way ANOVA followed by Šidak test).

### Postnatal cardiac endothelial cells promote atrial cardiomyocyte polyploidy

Crosstalk between myocytes and non-myocytes is essential for maintaining cardiac function. In particular, bidirectional communications between CMs and cardiac ECs, mediated by cardiokine and angiocrine signals, regulate multiple aspects of early heart development^30^, chamber morphogenesis^24,25,31,32^, and vascular development^33^. To investigate how non-myocytes contribute to chamber-specific CM polyploidy, we focused on cardiac ECs, the most abundant non-myocyte population in the heart^23^, comprising both vascular ECs and EdCs.

We first immunostained P7 atrial sections for ERG (Fig. 2A), an EC/EdC nuclear marker. EC/EdC quantification revealed that the RA contains a higher proportion of ECs/EdCs than the LA (Fig. 2B), as well as a higher EC/EdC density per unit area of the myocardium (Fig. 2C). Consistently, flow cytometry analysis showed a higher number of CD31+ EC/EdC per mg atrial tissue in the RA than in the LA (Supplementary Fig. 1A). To test whether atrial ECs (aECs) modulate aCM cytokinesis and polyploidy, we performed indirect co-culture experiments using conditioned medium. Primary rat aCMs were treated with conditioned medium collected from primary P7 rat aECs (Supplementary Fig. 1B) or atrial fibroblasts (aFbs). Cell cycle activity was assessed by EdU incorporation, whereas cytokinesis failure was quantified using an established EdU pulse-chase binucleation assay^20^. We observed that conditioned medium from aECs, but not aFbs, significantly increase both cell cycle activity (Fig. 2D) and cytokinesis failure (Fig. 2E) of primary LA and RA CMs, effects that were abolished by heat denaturation of conditioned medium (aEC_b) prior to treatment. Conditioned medium from human umbilical vein endothelial cells (HUVEC) and human cardiac microvascular endothelial cells (HCMEC) produced similar, although weaker, effects on aCM cell cycle activity (Supplementary Fig. 1C). Altogether, these data indicate that aECs can promote both cell cycle activity and cytokinesis failure in primary aCMs, likely through secreted angiocrine factors. However, because EC abundance is higher in the RA, EC number alone is unlikely to explain the chamber-specific pattern of aCM polyploidy. Instead, these observations point to qualitative differences in aEC-derived angiocrine factors between the LA and RA as a potential underlying mechanism.

**Figure 2.**
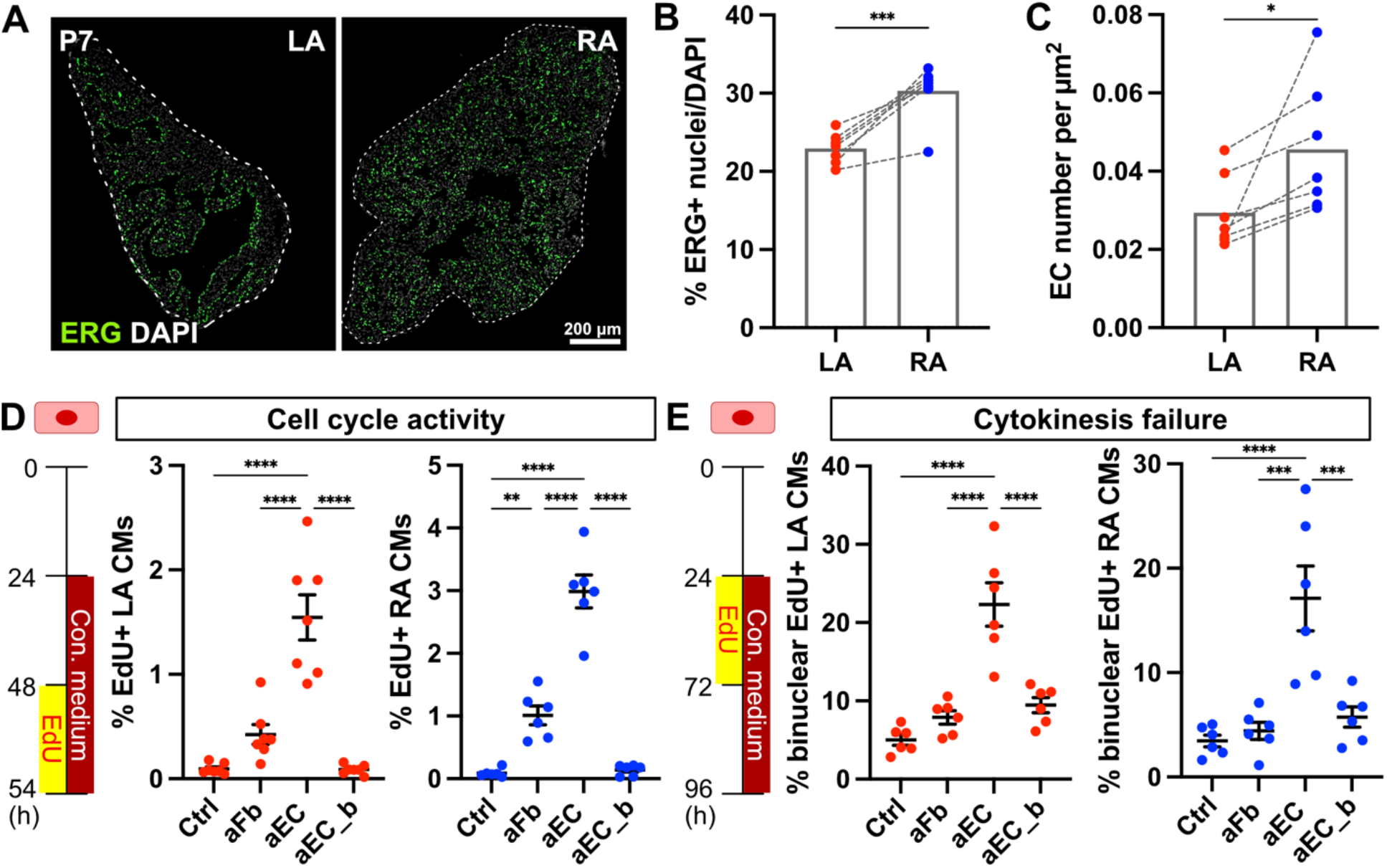
ECs/EdCs regulate aCM cell cycle entry and polyploidization via secreted factors. **(A)** Immunostaining for ERG (green), a nuclear marker for ECs/EdCs. Atrial sections were outlined by dashed lines. Scale bar represents 200 µm. **(B)** Quantification of the percentage of EC/EdC nuclei over total nuclei in LA and RA on P7 (n = 7) mice. LA and RA samples from the same mouse were connected with dashed line. **(C)** Quantification of EC/EdC density in LA and RA on P7 (n = 7). LA and RA samples from the same mouse were connected with dashed line. **(D)** Schematic of the EdU incorporation assay shown on the left. Quantification of the percentage of EdU+ LA (middle, n = 7) and RA CMs (right, n = 6) after treatment with normal or heat denatured (_b) conditioned medium collected from atrial fibroblasts (aFb) and atrial endothelial cells (aECs). **(E)** Schematic of the EdU pulse-chase assay shown on the left. Quantification of the percentage of EdU+ LA (middle, n = 6) and RA CMs (right, n = 6) after treatment with normal or heat denatured (_b) conditioned medium collected from aFbs and aECs. Values represent mean ± s.e.m. *B, C* (* *P < 0*.*0332, \**\*\* *P < 0*.*0002* by Two-tailed paired Student’s T test); *D, E* (*\**\*\* *P < 0*.*0002*, **** *P < 0*.*0001* by One-Way ANOVA followed by Tukey test).

### EDN1 as a candidate angiocrine factor that regulates atrial cardiomyocyte ploidy

To identify candidate angiocrine factors that regulate aCM polyploidy, we performed RNA-seq on P4 mouse LA and RA. We first selected chamber-enriched genes, defined by an absolute log2 foldchange (log2FC) >0.2, yielding 3752 genes. We then filtered this list for genes enriched in ECs based on a published single-cell RNA-seq dataset of the postnatal mouse heart^34^ (= 652 genes), and further narrowed these genes to known secreted factors (= 52; Fig. 3A). Among the top 10 LA-enriched EC/EdC-derived secreted factors was endothelin-1 (*Edn1*). In addition to its well-established role as a potent vasoconstrictor^35^, EDN1 has been implicated in several aspects of cardiac biology, including CM contractility^36,37^, terminal differentiation^38^, homeostasis and survival^39,40^. We therefore selected *Edn1* for further analysis.

To validate these findings, we re-analyzed the single-cell RNA-seq dataset of the postnatal mouse atria^34^, and found that *Edn1* was enriched in ECs compared with other non-myocyte populations (Supplementary Fig. 2A), with particularly high expression in EdCs (Fig. 3B). We next focused on genes encoding secreted factors within the vascular ECs and EdCs clusters and filtered for those showing significant chamber-enrichment specifically between P4 and P9, the developmental time window during which LA CM cytokinesis failure and binucleation increase substantially (Figs. 1B, D). Consistent with our bulk RNA-seq analysis, *Edn1* emerged as one of the top candidates, showing only modest LA enrichment at P1, but became more than two-fold enriched in the LA from P4 to P9 (Figs. 3C, D). We further validated these findings by qPCR using enriched mouse LA and RA ECs (Supplementary Fig. 2B). *Edn1* expression in LA ECs increased significantly from P1 to P7 before returning to P1 levels by P56, whereas *Edn1* expression in RA ECs remained largely unchanged over the same period (Fig. 3E). Accordingly, the relative enrichment of *Edn1* in LA versus RA ECs increased markedly from P1 to P7, rising from approximately two-fold to five-fold (Fig. 3E). This temporal pattern positively correlated with the divergent binucleation trajectories of LA and RA CMs during this period (Figs. 1B, D). We next examined the expression of the two G protein-coupled receptors for EDN1. Expression of endothelin receptor type a (*Ednra*), rather than type b (*Ednrb*), was highly enriched in aCMs (Fig. 3F) and was expressed at comparable levels between LA and RA CMs (Fig. 3G). Based on these findings, we hypothesized that EC/EdC-derived EDN1 inhibits CM cytokinesis and promotes polyploidy in the LA.

**Figure 3.**
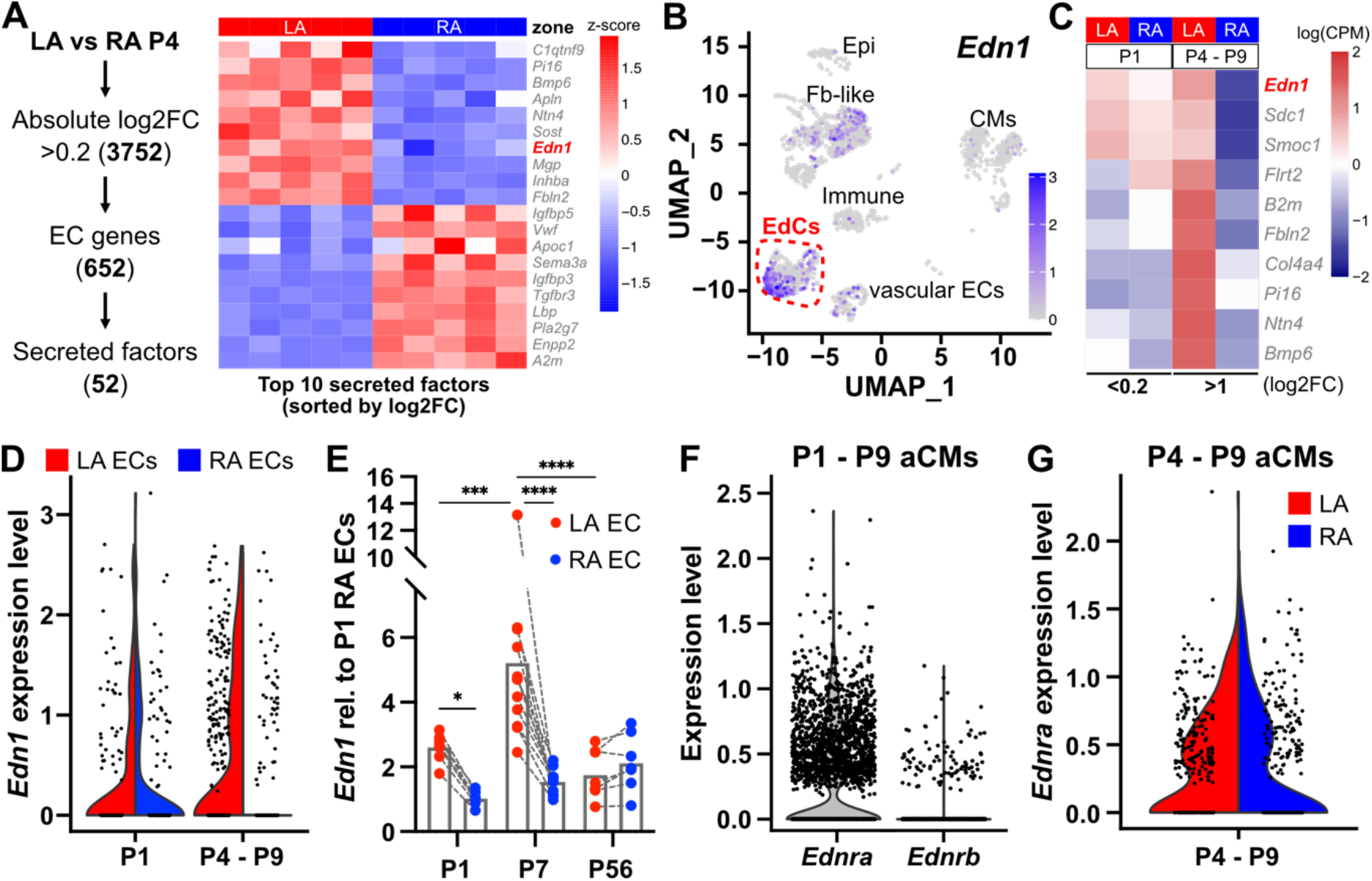
Identification of *Edn1* as a candidate angiocrine factor. **(A)** RNA-seq of P4 LA and RA; strategy for candidate gene filtering shown on the left. Heatmap for top 10 secreted factors from LA and RA shown on the right. **(B)** UMAP showing *Edn1* expression in mouse atria from P1 to P9. EdCs cluster indicated by red lines. Epi – Epicardial cells; Fb-like – Fibroblast-like cells; CMs – Cardiomyocytes. **(C)** Heatmap of secreted factors from P1 and P4 to P9 EdCs selected from *B*. **(D)** Violin plot of *Edn1* expression in P1 and P4 to P9 LA and RA ECs (i.e., EdCs and vascular ECs). **(E)** Quantification of *Edn1* expression in isolated LA and RA ECs/EdCs by qPCR at P1 (n = 7), P7 (n = 12), and P56 (n = 8). LA and RA samples from the same mouse were connected with dashed line. **(F)** Violin plot of *Ednra* and *Ednrb* expression in aCMs between P1 to P9. **(G)** Violin plot of *Ednra* expression in LA and RA CMs between P4 to P9. Values represent mean ± s.e.m. *E* (*** *P < 0*.*0002, \*\**\*\* *P < 0*.*0001* by Two-Way ANOVA followed by Tukey test).

### Endothelin-1 signaling promotes left atrial cardiomyocyte cytokinesis failure and polyploidy

To investigate the function of Edn1 signaling *in vivo*, we treated pups with Bosentan, a dual EDNRA/EDNRB antagonist, from P1 to P6 and assessed CM cytokinesis failure at P7 using the EdU pulse-chase assay described in Fig. 1D. No gross morphological defects or obvious changes in heart and body weight were observed in pups treated with Bosentan (Supplementary Fig. 3A). Bosentan treatment significantly reduced the proportion of binuclear EdU+ LA CMs (Fig. 4A), while the nuclear ploidy of mononuclear EdU+ CMs remained predominately diploid in both DMSO and Bosentan-treated groups (Fig. 4B). These findings suggest that inhibition of Edn1 signaling promotes successful cytokinesis in LA CMs. Notably, this effect was not observed in RA CMs from the same cohorts (Fig. 4A), or in vCMs (Fig. 4C), indicating a chamber-specific requirement for Edn1 signaling in LA CM cytokinesis failure.

Next, we asked whether EDNRB, a EdCs- and vascular ECs-enriched receptor (Supplementary Fig. 3B), also contributes to aCM polyploidy. Treatment with the EDNRB-specific antagonist BQ788 did not affect CM cytokinesis in both LA and RA (Fig. 4D), or in the ventricle (Fig. 4E). Altogether, these data suggest that Edn1 signaling inhibits cytokinesis specifically in LA CMs, most likely through EDNRA rather than EDNRB.

**Figure 4.**
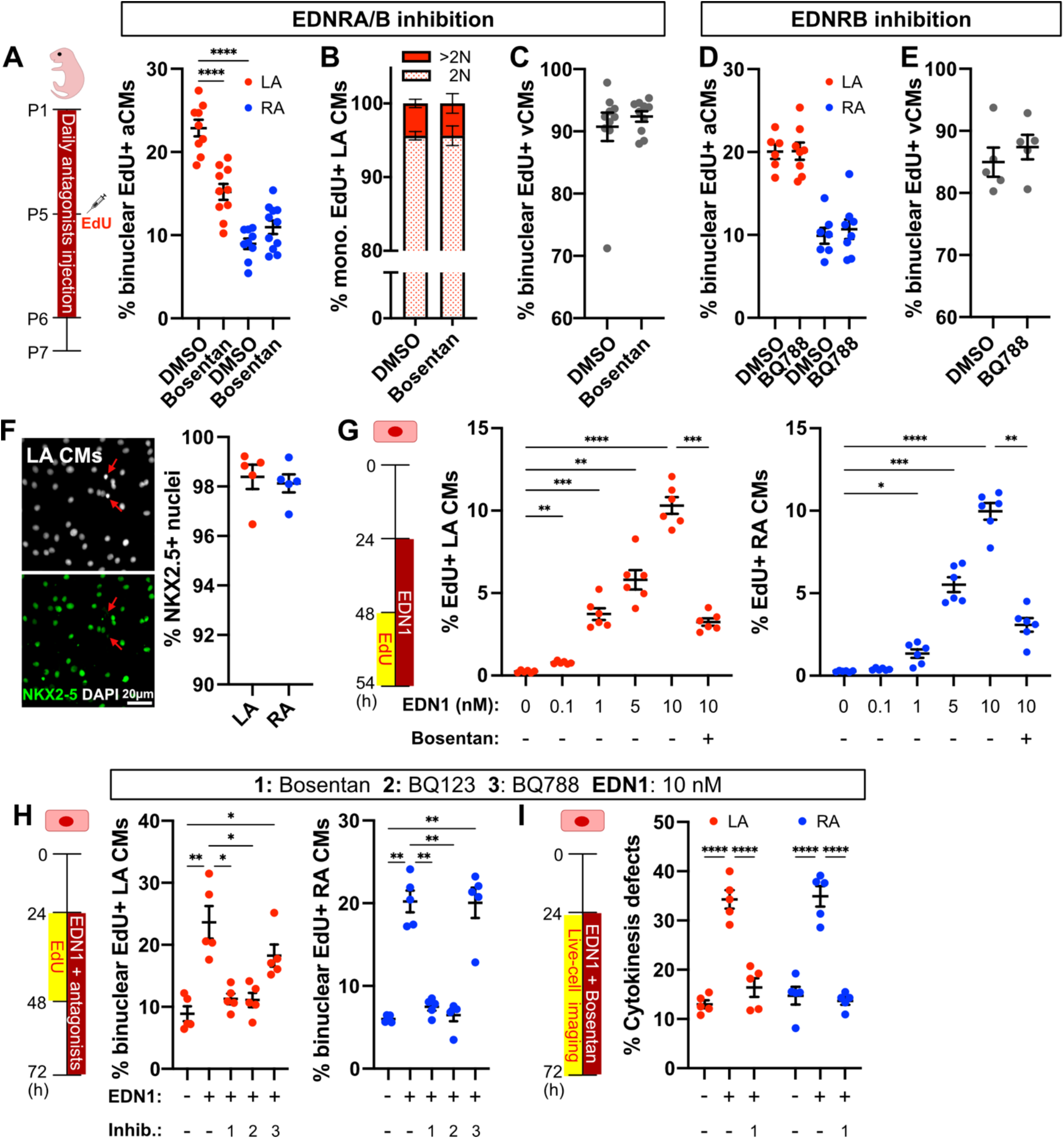
Edn1 signaling inhibits aCM cytokinesis *in vivo* and *in vitro*. **(A)** Schematic of the experimental design shown on the left. Quantification of the percentage of EdU+ binuclear LA and RA CMs at P7 in DMSO-(n = 9) and Bosentan-treated (n = 10) pups. **(B)** Quantification of nuclear ploidy of mononuclear EdU+ LA CMs (DMSO, n = 5; Bosentan, n = 6) from *A*. **(C)** Quantification of the percentage of EdU+ binuclear vCMs at P7 in DMSO-(n = 10) and Bosentan-treated (n = 10) pups. **(D)** Quantification of the percentage of EdU+ binuclear LA and RA CMs at P7 in DMSO-(n = 6) and BQ788-treated (n = 8) pups. **(E)** Quantification of the percentage of EdU+ binuclear vCMs at P7 in DMSO-(n = 5) and BQ788-treated (n = 5) pups. **(F)** Examples of primary LA CMs immunostained for NKX2-5, a nuclear CM marker, shown on the left. Quantification of the purity of aCM cultures (%NKX2-5+ nuclei, n = 5) shown on the right. Scale bar represents 20 µm. Non-myocytes indicated by red arrows. **(G)** Schematic of the EdU incorporation assay shown on the left. Quantification of the percentage of EdU+ primary LA (middle, n = 6) and RA (right, n = 6) CMs. **(H)** Schematic of the EdU pulse-chase binucleation assay shown on the left. Quantification of the percentage of EdU+ binuclear primary LA (middle, n = 5) and RA (right, n = 5) CMs. **(I)** Schematic of the experimental design shown on the left. Quantification of the percentage of cycling primary LA (middle, n = 5) and RA CMs (right, n = 5) undergo cytokinesis failure. Values represent mean ± s.e.m. *A, G, H* (* *P < 0*.*0332, \**\*\* *P < 0*.*0002*, **** *P < 0*.*0001* by One-Way ANOVA followed by Tukey test); *D* (*P > 0*.*05* by One-Way ANOVA followed by Tukey test); *C, E* (*P* > 0.05 by two-tailed Student’s t test); *I* (**** *P < 0*.*0001* by Two-Way ANOVA followed by Tukey test).

To test whether Edn1 signaling can act directly on aCMs, we used highly pure primary rat aCM cultures (∼98% purity; Fig. 4F). EDN1 peptide treatment promoted cell cycle entry in both LA and RA CMs in a dose-dependent manner, and this effect was blocked by Bosentan (Fig. 4G). Furthermore, EdU pulse-chase assays showed that EDN1 peptide was sufficient to induce cytokinesis failure in cycling primary LA and RA CMs (Fig. 4H). Consistent with our *in vivo* findings, these effects were abolished by Bosentan and by BQ123, an EDNRA-specific antagonist, but not by BQ788 (Fig. 4H). We further validated these observations by live cell imaging of primary LA and RA CMs (Fig. 4I). Altogether, these findings indicate that EDN1 can act directly on aCMs to promote cytokinesis failure through EDNRA-dependent signaling.

### Endothelin-1 signaling regulates cardiomyocyte polyploidy via Wnt signaling

To elucidate the molecular mechanisms by which Edn1 signaling regulates CM cytokinesis and polyploidy, we treated mouse pups with Bosentan and isolated LA and RA CMs for RNA-seq at P7 (Fig. 5A). All samples showed substantially higher expression of CM marker genes, including *Myl7* and *Nppa*, than markers of ECs (*Cdh5, Pecam1*), leukocytes (*Ptprc*), and fibroblasts (*Pdgfra*^41^, *Thy1*^42^; Supplementary Fig. 4), indicating that our isolation and purification protocol yielded enriched aCM populations. Consistent with the low level of *Edn1* expression in the RA (Fig. 3E), Bosentan treatment induced a markedly larger transcriptional response in LA CMs than in RA CMs, with 959 differentially expressed genes in LA CMs compared with only 27 in RA CMs using a cutoff of absolute log2FC >0.58 and padj <0.05 (Fig. 5B). We focused our analysis on genes that were highly enriched in LA CMs compared with RA CMs at baseline (LA ctrl versus RA ctrl: log2FC >2) and were downregulated after Bosentan treatment. Among the 304 genes, we identified several known Wnt inhibitors, including *Sfrp2, Sfrp4, Frzb*, and *Apcdd1* (Figs. 5C, D). *Sfrp2, Sfrp4*, and *Frzb* belong to the secreted Frizzled-related proteins (SFRPs) family due to their homology to the Wnt-binding Frizzled proteins^43^, while *Apcdd1* encodes a membrane-bound glycoprotein that functions upstream of ß-CATENIN to inhibit Wnt signaling^44^. Wnt signaling is a potent inducer of CM proliferation, including in human induced pluripotent stem cell-derived CMs^45^. Similarly, in postnatal mice, *Sfrp2*-mediated inhibition of canonical Wnt/ß-catenin signaling downstream of the transcription factor FOXO3 inhibits neonatal mouse CM proliferation and regeneration^46^. Altogether, we hypothesized that Edn1 signaling may maintain a pro-polyploidization CM state characterized by reduced Wnt pathway activity.

**Figure 5.**
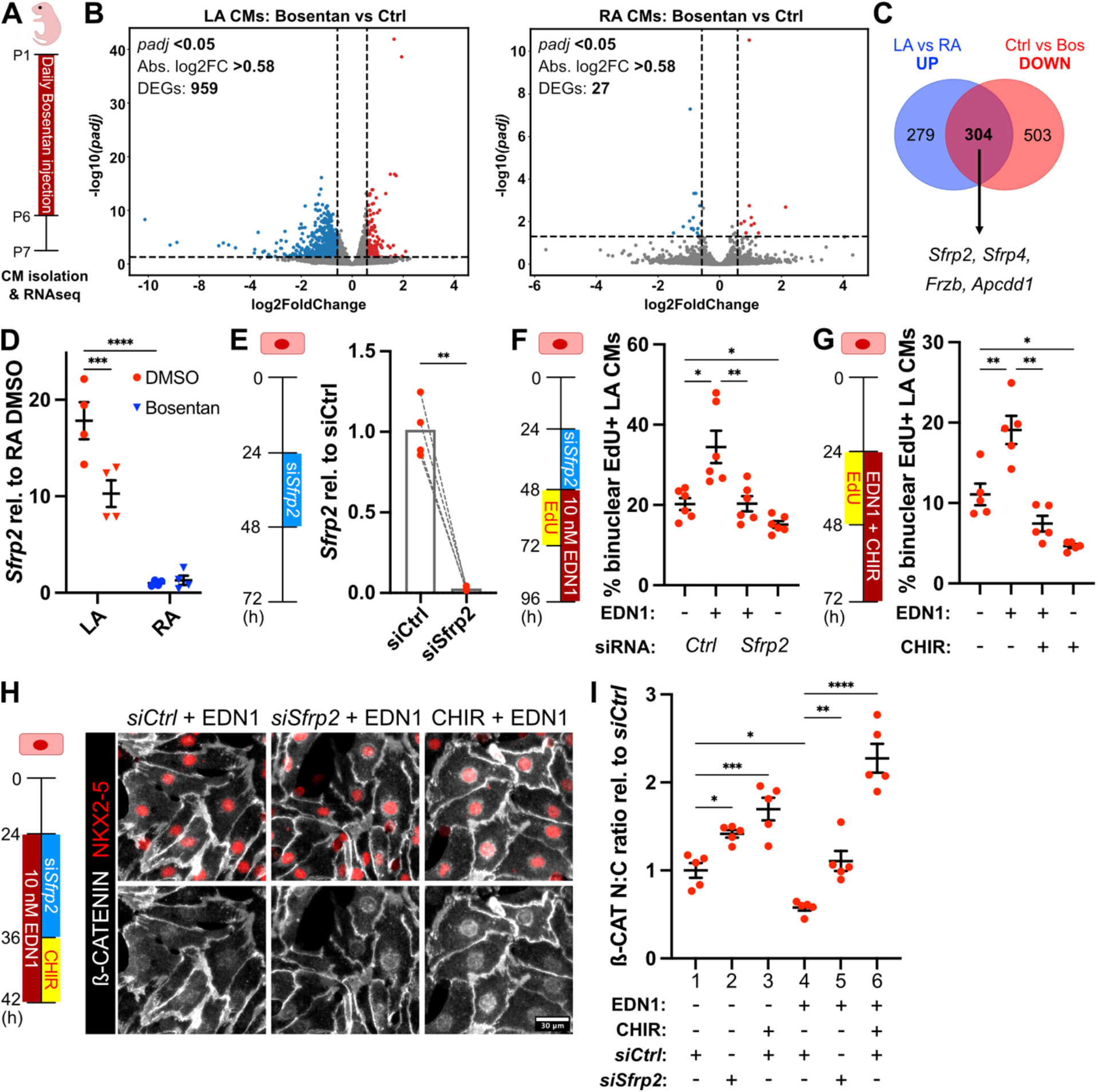
Edn1 signaling regulates aCM cytokinesis failure by suppressing Wnt signaling. **(A)** Schematic of the experimental design. **(B)** Volcano plots of gene expression in LA and RA CMs (Bosentan vs Ctrl). Up- and downregulated genes, defined as absolute log2FC>0.58 and padj <0.05, are denoted in red and blue, respectively. **(C)** Vann diagram between upregulated genes in LA CMs at baseline (blue) and downregulated genes in LA CMs after Bosentan treatment (red). **(D)** RNA-seq data of *Sfrp2* expression in LA and RA CMs from DMSO-(n = 4) and Bosentan-treated (n = 4) pups. **(E)** Schematic of the experimental design shown on the left. Quantification of *Sfrp2* expression by qPCR in primary LA CMs transfected with *siCtrl* and *siSfrp2* (n = 4). *siCtrl* and *siSfrp2* samples from the same litter were connected with dashed line. **(F)** Schematic of the EdU pulse-chase assay shown on the left. Quantification of the percentage of EdU+ binuclear primary LA CMs in *siCtrl*- and *siSfrp2*-transfected groups (n = 6). **(G)** Schematic of the EdU pulse-chase assay shown on the left. Quantification of the percentage of EdU+ binuclear primary LA CMs in control and CHIR-treated groups (n = 5). **(H)** Schematic of the experimental design shown on the left. Examples of primary LA CMs immunostained for NKX2-5 and ß-CATENIN shown on the right. Scale bar represents 30 µm. **(I)** Quantification of the nuclear to cytoplasmic ratio of ß-CATENIN in primary LA CMs after treatments (n = 5). Values represent mean ± s.e.m. *D* (*\**\*\* *P < 0*.*0002*, **** *P < 0*.*0001* by Two-Way ANOVA followed by Šidak test); *E* (** *P* < *0*.*0021* by two-tailed paired Student’s t test); *F, G, I* (* *P < 0*.*0332*, ** *P* < *0*.*0021, \**\*\* *P < 0*.*0002*, **** *P < 0*.*0001* by One-Way ANOVA followed by Šidak test).

To test this hypothesis, we performed loss- and gain-of-function experiments to modulate Wnt activity in primary LA CMs. Among the four Wnt inhibitors identified, we focused on *Sfrp2* because of its previously reported role in regulating mouse CM proliferation, and examined its function using siRNAs (Fig. 5E). *Sfrp2* knockdown promoted successful cytokinesis (*siCtrl* versus *siSfrp2*) and rescued the cytokinesis failure induced by EDN1 treatment (*siCtrl* + EDN1 versus *siSfrp2* + EDN1) in primary LA CMs (Fig. 5F). Next, we activated Wnt signaling with CHIR, a GSK-3ß inhibitor, which promoted successful cytokinesis (CHIR – versus CHIR +) and rescued EDN1-induced cytokinesis failure (EDN1 versus EDN1 + CHIR) in primary LA CMs (Fig. 5G). We next assessed Wnt activity in primary LA CMs by quantifying the nuclear-to-cytoplasmic ratio of ß-CATENIN (Fig. 5H). Compared with controls, both *Sfrp2* knockdown and CHIR increased the nuclear-to-cytoplasmic ß-CATENIN ratio in primary LA CMs, indicating increased Wnt activity (column 1 versus columns 2 and 3; Fig. 5I). Conversely, EDN1 peptide treatment significantly reduced Wnt activity (column 1 versus column 4; Fig. 5I), consistent with its effect on cytokinesis failure. Importantly, this reduction in Wnt activity was rescued by either *Sfrp2* knockdown or CHIR treatment (column 4 versus columns 5 and 6; Fig. 5I). Altogether, these findings suggest that Edn1 signaling promotes LA CM cytokinesis failure and polyploidization, at least in part, through modulating Wnt pathway activity.

## Discussion

CM polyploidy, arising from endoreplication or endomitosis, is a hallmark of CM maturation in mice^47^ and human induced pluripotent stem cell-derived CMs^48^. The postnatal transition from a diploid to a polyploid state is associated with CM cell cycle exit and the loss of endogenous regenerative capacity^6-8^. Although it remains unclear whether increasing the abundance of mononuclear diploid CMs alone is sufficient to promote cardiac regeneration^49-52^, strategies that enhance adult CM cell cycle entry and cytokinesis improve cardiac function after MI^12,53,54^. Despite the importance of this process during heart development and regeneration, the mechanisms that drive CM polyploidization remain poorly understood. Here, we show that both the timing and extent of postnatal mouse CM polyploidization are highly chamber-specific, and identify cardiac ECs/EdCs as regulators of LA CM cytokinesis, at least in part through Edn1 signaling (Fig. 6).

**Figure 6.**
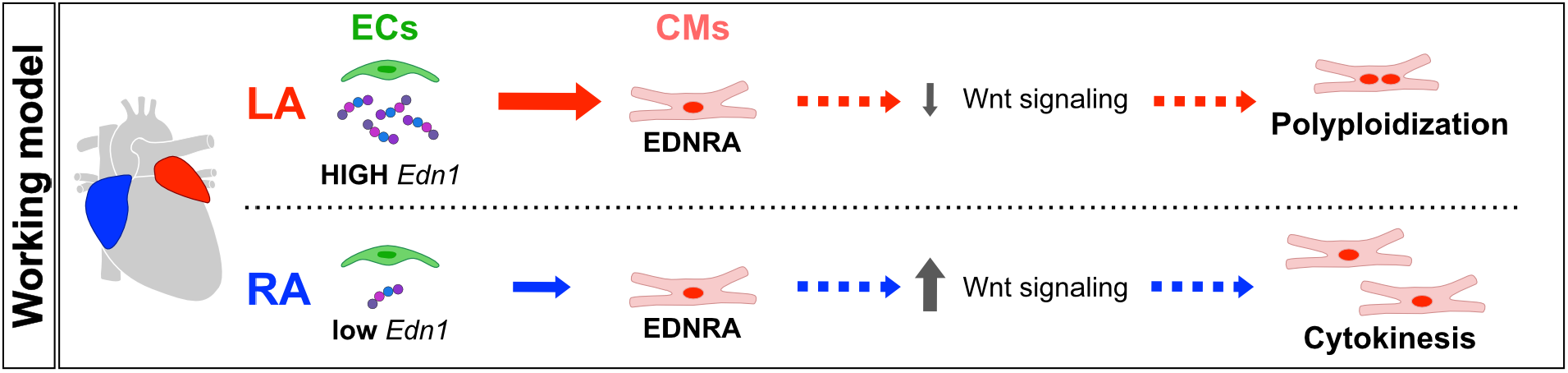
Working model.

vCM polyploidization has been extensively characterized through comparative studies across species^1,55^, mouse strains^8,29^, and developmental stages^5^, whereas much less is known about aCM polyploidy. Adult aCMs in mice^14^ and birds^56^ have lower levels of multinucleation and/or nuclear ploidy than their ventricular counterparts. Our time-course analysis further revealed that LA CMs acquire a significantly higher level of binucleation than RA CMs, reaching ∼25% in the LA compared with ∼12% in the RA by 8 weeks of age. This chamber difference emerged around P7 and correlates with the higher frequency of cytokinesis failure among cycling LA CMs than RA CMs (∼20% versus ∼10%; Fig. 1D), although potential differences in baseline cell cycle activity may also contribute to the total number of polyploid CMs. Intriguingly, the timing and the magnitude of chamber-specific aCM polyploidy appear to differ across species. In rats, LA CMs showed higher levels of binucleation than RA CMs mainly at adult stages^16^, whereas adult birds display a similar but less pronounced LA–RA difference^56^. A direct comparison of nucleation between human LA and RA CMs is still lacking. Further studies are therefore required to determine whether chamber-specific CM polyploidy is conserved across mammals and how it relates to chamber- and species-specific physiology.

The majority of mechanistic studies of CM polyploidy have focused on CM-intrinsic regulators^18^. Dysregulation of cell cycle and cytokinesis regulators during development, including *Ccng1*^57^, *Ect2*^58^, and mislocalization of ANILIN^59,60^, have been shown to alter CM ploidy. Beyond core cell cycle regulators, a number of CM-autonomous pathways have also been implicated in CM cytokinesis and ploidy control, including E2F transcription factor family members *E2f7* and *E2f8*^50^, the CM-specific kinase *Tnni3k*^8^ (although its exact role remains controversial^10^), *Rbpms*^61^ *Lmnb2*^62^, PIDDosome^63^, p38 MAPK^59,64^, NRG1^65^ and its co-receptor ERBB2^66^, thyroid hormone signaling^1,67^, and the Hippo pathway^68-70^. In contrast, much less is known about how non-myocytes regulate CM polyploidization. We and others have previously shown that Periostin-expressing cardiac fibroblasts^19^ and the postnatal cardiac extracellular matrix^20^ promote vCM cytokinesis failure and binucleation. Here, we extend this concept to aCMs and identify cardiac ECs/EdCs as important regulators of chamber-specific aCM polyploidization. Among the candidate angiocrine factors identified in our analysis, EDN1 emerged as a compelling LA-enriched EC/EdC-derived signal, which has been implicated in multiple aspects of CM biology. In addition, EDN1 is upregulated in the LA appendage of patients with atrial fibrillation^71^, a disease associated with structural remodeling and increased CM nuclear ploidy^15^. Although correlative, these human observations suggest that Edn1 signaling may have a broader relevance in atrial remodeling under pathological conditions. Our data suggest that Edn1 signaling promotes LA CM cytokinesis failure primarily through EDNRA. Pharmacological inhibition of Edn1 signaling with Bosentan reduced cytokinesis failure (Fig. 4A) and elicited a much stronger transcriptional response in LA CMs than in RA CMs (Fig. 5B). Given that LA and RA CMs expressed comparable levels of *Ednra* (Fig. 3G), and responded similarly to EDN1 peptide *in vitro* (Fig. 4H), differential *Edn1* availability between LA and RA (Fig. 3E) is likely the major contributor to the chamber-specific effect of Bosentan *in vivo*. However, what drives the LA-RA difference in *Edn1* expression in ECs/EdCs is currently unclear.

Several limitations in our study should be noted. First, although our expression analyses identify ECs/EdCs as a major source of *Edn1* in postnatal mouse atria (Supplementary Fig. 2A), pharmacological inhibition with Bosentan does not necessarily prove that EC/EdCs-derived EDN1 is the relevant ligand source *in vivo*; *Edn1* contributed from other non-myocyte populations, including fibroblasts, cannot be formally excluded (Fig. 3B). Secondly, EDN1 is known to act on multiple cardiac cell types, including fibroblasts^72,73^ and smooth muscle cells^35^. Hence, while our highly pure primary aCM culture experiments support a direct, CM-autonomous effect of EDN1 on cytokinesis failure (Figs. 4G, H), we cannot exclude potential contributions from alternations in fibroblast activity and blood pressure or other hemodynamic parameters after Bosentan treatment *in vivo*. Future EC/EdC-specific *Edn1* loss-of-function and CM-specific *Ednra* loss-of-function studies will be essential to identify the important cellular source of EDN1 and definitively establish the cell-autonomous function of Edn1 signaling in LA CM polyploidization.

Bi-directional communication between ECs and CMs has been implicated in various aspects of cardiac biology. During embryonic heart development, endocardial Notch signaling and NRG1 regulate trabeculation in the ventricle^24,25,74^, a signaling axis that is conserved in zebrafish^31,75^. In the atria, CM-derived ANGPT1 regulates ADAMTS expression in EdCs, thereby promoting cardiac jelly degradation and proper chamber morphogenesis^32^. In the adult mouse heart, *Vegfb* overexpression promotes angiogenesis-induced CM hypertrophy via VEGFR2 signaling in ECs, which can be attributed to increased NRG1 release from ECs^76,77^. The findings presented here highlight the previously unrecognized role for EdCs in postnatal mouse heart development by regulating chamber-specific CM polyploidization through differential Edn1 signaling.

## Methods

### Mice

All animal experiments were approved by the governmental (Regierungspräsidium Karlsruhe, Germany) and/or institutional committees, and were performed in accordance with guidelines for the care and ethical use of laboratory animals. C57BL/6JRj mice were purchased from Janvier Laboratories. All mice were housed on a 12h light/12h dark cycle with free access to food and drinking water.

### Animal experiments

5-Ethynyl-2’-Deoxyuridine (EdU, Baseclick, BCN-001), dissolved in PBS, was injected intraperitoneally at 50 mg/kg two days before heart harvesting. Bosentan (Merck, SML1265), dissolved in DMSO, was injected intraperitoneally at 4 mg/kg; same amount of DMSO was injected as controls. BQ788 (Biotechne, Cat. No. 1500), dissolved in DMSO, was injected intraperitoneally at 1 mg/kg; same amount of DMSO was injected as controls.

### Mouse cardiomyocyte dissociation

After removal, hearts were fixed with 4% paraformaldehyde (PFA, Sigma, P6148) at 4°C for 48 h. After washing with PBS, atria and ventricles were dissected and incubated in 50% w/v potassium hydroxide solution overnight at 4°C. After washing with PBS, tissue was transferred to a 100 μm cell strainer on a petri dish containing PBS and cells were released by smashing gently with the stamp of a 5- or 10-ml syringe. Single cell suspension was then spotted on a glass slide and airdried overnight.

### Mouse cardiac endothelial cells isolation

Left and right atria from P7 mice were dissociated using the Neonatal Heart Dissociation Kit (Miltenyi Biotec, 130-098-373) according to the manufacturer’s protocol. Afterwards, endothelial cells were purified by positive selection using CD31 microbeads (Miltenyi Biotec, 130-097-418) according to the manufacturer’s protocol.

### Rat atrial cardiomyocyte isolation and culture

Neonatal (P1 or P2) rats were sacrificed by decapitation, and the hearts were harvested. Left and right atria were then dissociated using the Neonatal heart dissociation kit (Miltenyi Biotec, 130-098-373) according to manufacturer’s protocol. Afterwards, CMs were purified using the Neonatal CM isolation kit (Miltenyi Biotec, 130-105-420) according to manufacturer’s protocol. LA and RA CMs (2 × 10^5^ cells/cm^2^) were plated on Phenoplate (PerkinELmer, 6055300) coated with 100 μg/mL fibronectin (Sigma, 341631). CMs were cultured in neonatal CM medium (DMEM/F12, GlutaMAX with 3 mM Na-Pyruvate, 0.1 mM Ascorbic acid, Insulin/transferrin/Na-Selenite solution (1:200), 0.2% BSA and 1X Pen/Strep) containing 10% FBS (Sigma, S0615). To perform the EdU pulse-chase assay, CMs were labeled with 20 μM EdU for 24 h, followed by two washes with neonatal CM medium to remove EdU and a 24 h chase. Cytokinesis defects were assessed by quantifying the percentage of binuclear EdU+ CMs. Each independent CM isolation was considered a separate biological replicate.

EDN1 peptide (PeptaNova, 4198-v), dissolved in 0.1% acetic acid, was treated at 10 nM; equal amount of acetic acid was used as controls. Bosentan (Merck, SML1265), BQ123 (Selleck Chemicals, S7883), and BQ788 (Biotechne, Cat. No. 1500) were treated at 10 μM, 20 μM, and 10 μM, respectively.

### siRNA transfection and CHIR treatment

Primary aCMs were transfected with 50 nM ON-TARGETplus SMARTpool siRNA targeting rat *Sfrp2* (Horizon Discovery; L2-088918-01-0005) or ON-TARGETplus Non-targeting Pool (D-001810-10-05) using Lipofectamine RNAiMAX (Thermo Fisher Scientific, 13778030) according to the manufacturers’ instructions. Cells were harvested 48 h after transfection for downstream analyses. 5 μM CHIR99021 (Sigma, SML1046) was used for Wnt signaling activation in aCMs.

### Live cell imaging of aCMs

Primary P2 rat aCMs were isolated and allowed to attach to culture plates for 24 h prior to treatment. Cells were then treated with 10 nM EDN1 and/or 10 μM Bosentan for 24 h. Following EDN1 treatment, nuclei were labeled with siR-DNA (Spirochrome, #SC007; 1:1000 dilution) in neonatal CM medium containing 10% FBS. Time-lapse imaging was performed using an Invitrogen™ EVOS™ M7000 Imaging System under standard cell culture conditions (37°C, 5% CO_2_). Single-plane images were acquired every 20 min over a 24 h period. CM cytokinesis was analyzed manually using Fiji.

### Rat atrial cardiac endothelial cells and fibroblasts isolation, culture and conditioned medium collection

Left and right atria from P7 rats were dissociated using the Neonatal heart dissociation kit (Miltenyi Biotec, 130-098-373) according to manufacturer’s protocol. Afterwards, endothelial cells were purified by positive selection using Dynabeads Pan Mouse IgG (Thermo Fisher Scientific, 11-041) conjugated with purified mouse anti-rat CD31 antibody (BD, 550300). The beads and antibody were incubated overnight at 4°C for conjugation prior to use. Fibroblasts were collected from the remaining cells by pre-plating on 10 cm Petri dishes for 1 h. Purified endothelial cells were cultured in endothelial cell growth medium MV2 (Promocell, #C-22022) and maintained in 10 cm Petri dishes (Sarstedt, 83.3902) coated with 1% gelatin (Sigma, G1890). Fibroblasts were cultured in DMEM/F12, GlutaMAX containing 10% FBS and 1X Pen/Strep. Conditioned medium was collected from atrial endothelial cells and fibroblasts at ∼80% confluency daily for three consecutive days; fresh medium was replenished after each collection. Collected medium were centrifuged at 1000 × g for 10 min, filtered through a 0.2 μm filter, aliquoted, and stored at −20°C. For experiments, conditioned medium was diluted 1:2 in MV2 supplemented with 10% FBS. Heat-inactivated controls were generated by heating conditioned medium at 95°C for 15 min.

### Immunostaining

For dissociated mouse CMs, cells were rehydrated with PBS and permeabilized with 0.5% Triton-X 100/PBS. After blocking (10% donkey serum/PBST, Sigma, S30) for at least 1 h, slides were incubated with primary antibodies at 4°C overnight. Secondary antibodies (Alexa fluor dyes, Thermo Fisher Scientific) were diluted (1:500) in blocking buffer and incubated for 1 h at room temperature together with DAPI (Sigma, D9542). EdU detection was performed using the Click-iT EdU Cell Proliferation Kit (Thermo Fisher Scientific, C10640) or Sensitive EdU Cell Proliferation Assay for Imaging (Baseclick, BCK-EDUPRO647IM100) according to manufacturer’s protocol. Samples were mounted using VECTASHIELD Vibrance Antifade Mounting Medium (Thermo Fisher Scientific).

For rat aCMs, cells were fixed with 4% PFA on ice and immunostaining was performed as described above. Samples were mounted with ibidi Mounting Medium with DAPI (ibidi, 50011-4).

Images were captured with a THUNDER Imager (Leica). Primary antibodies used are anti-NKX2.5 (1:300, R&D, AF2444), anti-N Cadherin (1:1500, Abcam, ab18203), anti-α-Actinin (1:500, Sigma, A7811), anti-Myl7 (1:1000, Proteintech, 17283-1-AP), anti-ERG (1:300, Abcam, EPR3864), and anti-ß-Catenin (1:500, BD, 610153).

### Quantification

All data quantification were done blindly except nuclear ploidy analysis. Quantification of nucleation of dissociated CMs was performed manually with Fiji; at least 100 to 150 EdU+ CMs were quantified per samples. To quantify the nuclear ploidy of dissociated CMs, integrated DAPI intensity of all non-myocytes on a single tile was measured automatically with a CellProfiler pipeline. Afterwards, the integrated DAPI intensity of each mononuclear EdU+ CMs from individual tile were measured manually with Fiji. The obtained value was then divided by the average integrated DAPI intensity of all non-myocytes from the corresponding tile. Ratio > 1.5 was considered polyploid while ratio < 1.5 was considered diploid; at least 80 to 100 EdU+ mononuclear CMs were quantified per samples. Quantification of nucleation of primary rat aCMs was performed manually with Fiji; at least 100 EdU+ CMs were quantified per samples. For quantification of the nuclear to cytoplasmic ratio of ß-CATENIN, fluorescence images were acquired using identical imaging settings across all groups. Several small regions of interest, in which single CMs can be clearly distinguished, were randomly selected and cropped for analysis; only mononuclear cells were included in the analysis. For each cell, nuclear ß-CATENIN was quantified by manually outlining NKX2-5+ nuclei, while cytoplasmic β-CATENIN signal was calculated by subtracting nuclear β-CATENIN signal from that in the whole cell (i.e., measured by outlining the cell boundary excluding membrane-associated signal).

The proportion of ECs was determined as the percentage of ERG-positive nuclei (counterstained with DAPI). All DAPI+ and ERG+ nuclei were automatically quantified in Fiji for each section. EC density was defined as the number of ERG-positive nuclei normalized to the atrial area (number/μm^2^).

### Quantitative PCR

Quantitative real-time PCR (qPCR) was performed using TaqMan probe–based assays. Total RNA was extracted using the RNeasy Mini Kit (Qiagen, 74106), and cDNA was synthesized using the QuantiTect Reverse Transcription Kit (Qiagen, 205313) according to the manufacturer’s instructions. qPCR reactions were carried out using TaqMan™ Fast Advanced Master Mix (Thermo Fisher Scientific, 4444557), *Edn1* TaqMan probes (Mm00438659_m1), and *Sfrp2* TaqMan probes (RN01458837_m1). Relative *Edn1* and *Sfrp2* expression were calculated using the ΔΔCt method and normalized to *Actb* (Mm02619580_g1) and *Rpl4* (Rn06291926_g1), respectively.

### Statistics

GraphPad Prism was used for statistical tests. Values represent mean ± s.e.m. Student’s t-test was used to compare between two groups. One-way or Two-way ANOVA were used for comparison between multiple groups.

### Tissue dissociation and flow cytometry

Hearts were harvested and dissected into left atrium (LA), right atrium (RA), and ventricle (V). Tissue weight was recorded prior to mechanical mincing into small fragments. Samples were then enzymatically digested in Dulbecco’s Modified Eagle Medium (DMEM) containing Liberase TM (Sigma, 5401127001) and DNase (Sigma, 10104159001) for 30 min at 37°C, with repeated mechanical dissociation using a 19-gauge needle.

Following digestion, cell suspensions were filtered through a 70 µm cell strainer, centrifuged at 400 × g for 5 min, and washed with FACS buffer (phosphate-buffered saline supplemented with 3% fetal bovine serum). Red blood cells were lysed using ACK (ammonium–chloride– potassium) lysis buffer for 2 min at room temperature. Cells were subsequently incubated with anti-mouse CD16/CD32 antibody for Fc receptor blocking (10 min, 4 °C), followed by staining with fluorochrome-conjugated antibodies diluted in FACS buffer for 20 min on ice. FxCycle dye was used for discrimination of live and dead cells. Flow cytometric analysis was performed using a BD FACS Canto II, and data were analyzed with FlowJo software.

### Bulk RNA-seq preprocessing and analysis

Bulk RNA-seq was performed on whole LA and RA tissue from P4 mice (n = 5). The raw sequence reads were aligned to the mouse reference genome mm10 (Ensembl GRCm38, v.2.7.4a) using STAR aligner (v. 2.7.11a). The sample-wise read files were aggregated into a count matrix. Minimal pre-filtering was applied by removing lowly expressed genes with summed counts across all samples of less than 10. Differential expression analysis between left and right atria was computed using DESeq2 (v.3.18) and significantly differentially expressed genes were identified based on an absolute log2 fold-change (abs. log2FC) ≥ 0.2 and an adjusted p-value (p.adj) cutoff of 0.05.

To identify EC-enriched factors, we utilized a publicly available scRNA-seq dataset (GSE193346). The data was processed using the Seurat package (v4.4.0). The dataset was subset to include only atrial cells from postnatal time points P4 to P9. EC-enriched markers were identified using Seurat’s FindAllMarkers function with default thresholds, specifically selecting genes significantly enriched in endocardial and vascular EC clusters. These genes were then intersected with the bulk RNA-seq differentially regulated gene list. The resulting gene list was further filtered for secreted factors using the Secreted Protein Database (SEPDB, Mouse), retaining only those annotated as “Secreted” or “Extracellular” in the “Subcellular Location” column.

### Single cell RNA-seq analysis

UMAP plots were generated in Seurat to visualize gene expression within the atrial P4 to P9 subset of GSE193346. For the heatmap, genes were filtered to identify those with dynamic postnatal expression. Pseudo-bulk profiles were generated by computing the expression average per timepoint and zone (pseudo-bulk abs. log2FC ≤ 0.2 or p.adj ≥ 0.05 at P1, but abs. log2FC ≥ 1 and p.adj ≤ 0.05 between P4 to P9).

Violin plots for gene and receptor expression were created using ggplot2. To ensure unbiased comparison between chambers, cell numbers were downsampled to equal counts per timepoint for LA and RA ECs. For receptor analysis (Panel F), expression was analyzed specifically within the aCM cluster across the P1 to P9 developmental window.

## Data availability

Bulk RNA-seq data will be deposited to GEO and publicly available (accession no. pending).

## Acknowledgements

We acknowledge the data storage service SDS@hd at Heidelberg University (INST 35/1314-1 FUGG and INST 35/1503-1 FUGG) and are grateful for the support from the animal facilities of the Medical Faculty Mannheim, Heidelberg University. This work was supported by funding from the DFG within CRC1366 ‘Vascular control of organ function’ (project number 394046768) to CCW, MS, and the Emmy Noether Programme (project number 521708989) to CCW. This study was supported through state funds approved by the State Parliament of Baden-Württemberg for the Innovation Campus Health + Life Science Alliance Heidelberg Mannheim.

## Author Contributions

Project conceptualization, experimental design, funding acquisition: C-C.W., T.L. Data acquisition: T.L, S.K., Y.A, E.Z. Data analysis: C-C. W., T.L, S.K. Y.A., L.R., M.S. Manuscript writing: C-C.W., T.L, S.K., Y.A, L.R.

