## Supplementary Figures for "Endothelin-1 signaling regulates chamber-specific mouse atrial cardiomyocyte cytokinesis and polyploidy"

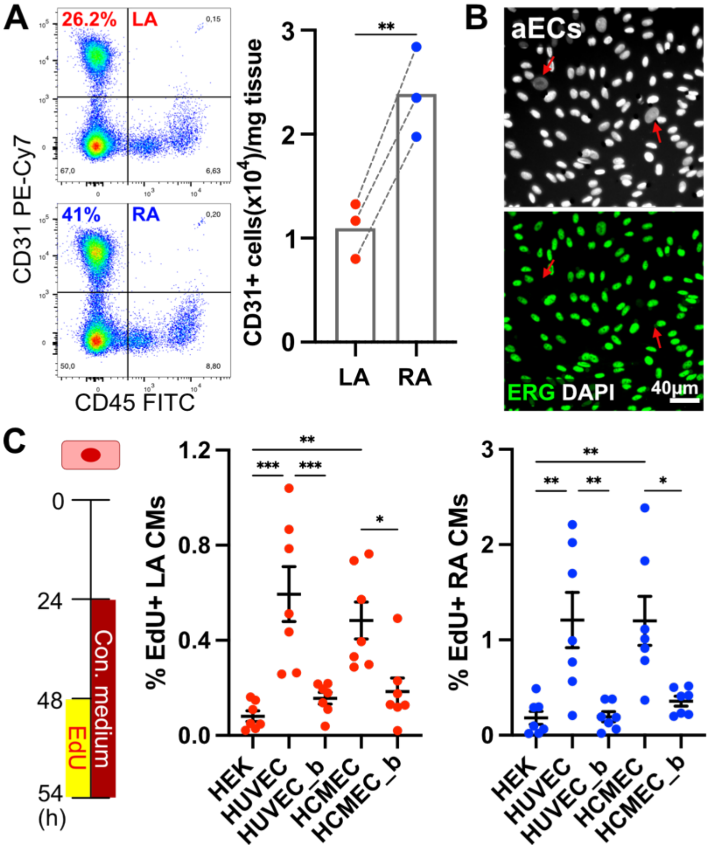


**Supplementary Figure 1.** **(A)** Representative flow cytometry plots shown on the left. Quantification of the number of CD31+ ECs/EdCs per mg tissue (n = 3). LA and RA from the same mouse was connected with dashed line. **(B)** Examples of primary aECs isolated from P7 rats immunostained for ERG (nuclear marker for ECs). Scale bar represents 40 μm. Non-ECs indicated by red arrows. **(C)** Schematic of the EdU incorporation assay shown on the left. Quantification of the percentage of EdU+ LA (middle, n = 7) and RA (right, n =7) CMs after treatment with normal or heat denatured (_b) conditioned medium collected from HEK293 cells (HEK), HUVECs, and HCMECs. Values represent mean ± s.e.m. *A* (** *P < 0.0021* by two-tailed paired Student's t test); *C* (* *P < 0.0332,* ** *P < 0.0021,* *** *P < 0.0002* by One-Way ANOVA followed by Tukey test).

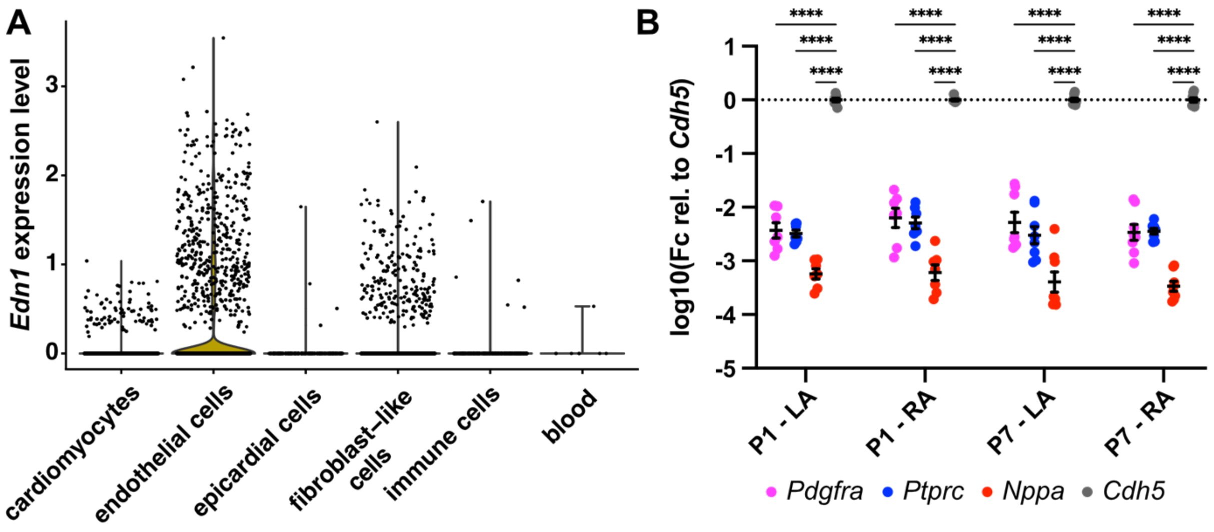


**Supplementary Figure 2.** **(A)** Violin plot of *Edn1* expression in different cell clusters in atria of P1 to P9 mice. **(B)** Quantification of marker genes expression (Fb: *Pdgfra;* Immune: *Ptprc*; CM: *Nppa*) relative to *Cdh5* (EC marker) in isolated primary LA and RA ECs using qPCR. Values represent mean ± s.e.m. *B* (**** $P < 0.0001$ by Two-Way ANOVA followed by Dunnett test).

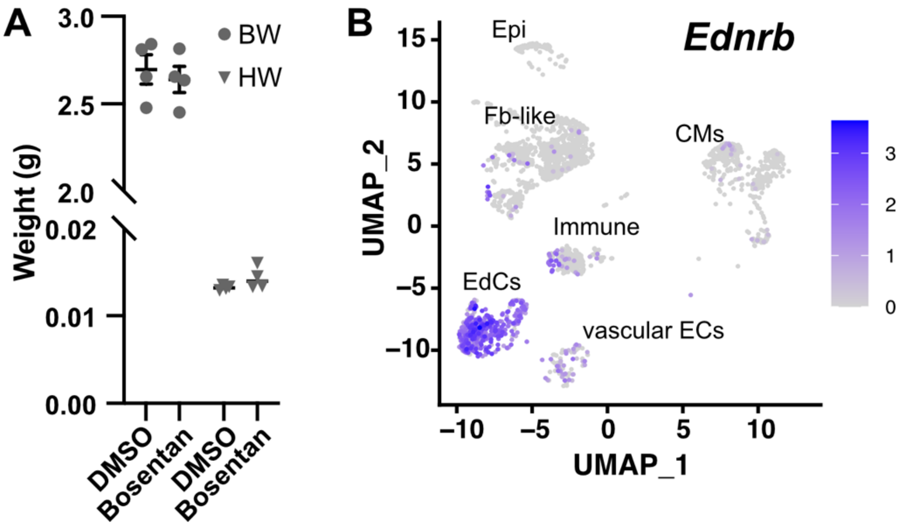


**Supplementary Figure 3. (A)** Quantification of body weight and heart weight of P7 mice after Bosentan treatment (n = 4). **(B)** UMAP plot of *Ednrb* expression in atria from P1 to P9 mice. Values represent mean ± s.e.m. *A* ($P > 0.05$ by Two-tailed Student's t test).

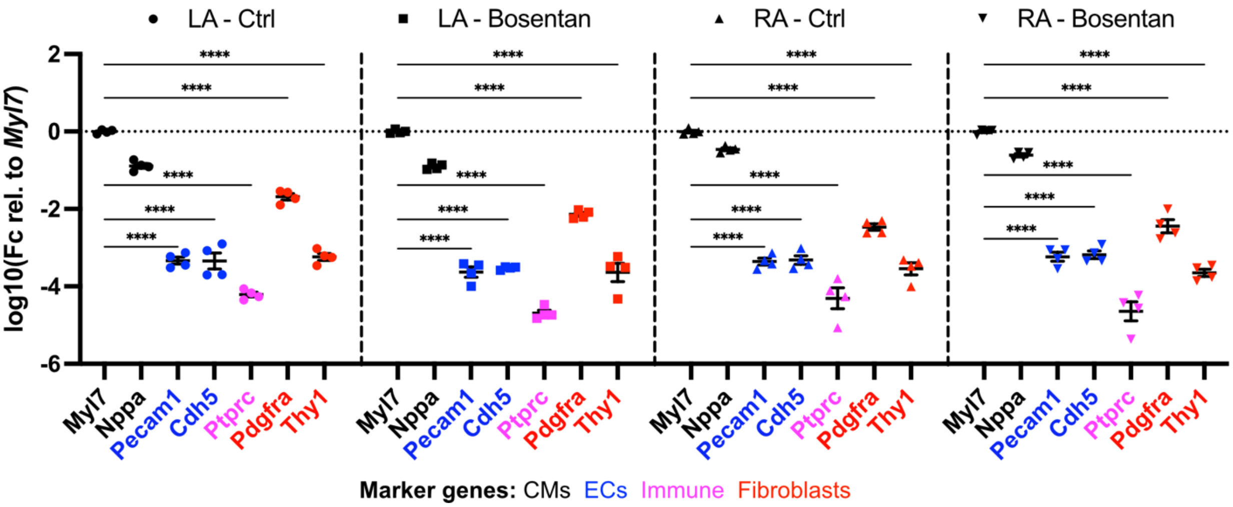


**Supplementary Figure 4.** Analysis of marker genes expression relative to *Myl7* (aCM marker) from RNA-seq data of enriched LA and RA CMs (n = 4) isolated from control- and Bosentan-treated groups. Values represent mean ± s.e.m. (**** $P < 0.0001$ by Two-Way ANOVA followed by Dunnett test).
